# Sapling: Inferring and Summarizing Tumor Phylogenies from Bulk Data using Backbone Trees

**DOI:** 10.1101/2024.04.10.588891

**Authors:** Yuanyuan Qi, Mohammed El-Kebir

## Abstract

Cancer phylogenies are key to understanding tumor evolution. There exists many important downstream analyses that takes as input a single or small number of trees. However, due to uncertainty, one typically infers many, equally-plausible phylogenies from bulk DNA sequencing data of tumors. We introduce Sapling, a heuristic method to solve the Backbone Tree Inference from Reads problem, which seeks a small set of backbone trees on a smaller subset of mutations that collectively summarize the entire solution space. Sapling also includes a greedy algorithm to solve the Backbone Tree Expansion from Reads problem, which aims to expand an inferred backbone tree into a full tree. We prove that both problems are NP-hard. On simulated and real data, we demonstrate that Sapling is capable of inferring high-quality backbone trees that adequately summarize the solution space and that can expanded into full trees.

## Introduction

Cancer results from an evolutionary process during which somatic mutations accumulate in a population of cells [23]. This process results in intra-tumor heterogeneity, i.e. the presence of multiple clones with distinct sets of mutations, with important implications on cancer treatment [20]. Researchers model cancer evolution with a *phylogeny*, which is rooted tree whose nodes correspond to clones. These trees are used in several downstream analysis [26]. These downstream analyses typically require a single or a small number of phylogenies per patient. However, deconvolution of bulk DNA measurements may lead one to infer a large solution space of equally-plausible phylogenies [25].

There are three classes of methods that attempt to overcome this mismatch between the existence of large solution spaces and downstream analysis requirements. First, several approaches attempt to sample a small number of high-likelihood trees [18, 19, 29, 5]. Second, there exist several methods that attempt to summarize a given solution space of trees with one or more consensus trees [12, 6, 11, 16, 9, 1, 10]. Third, there exist approaches that use repeated evolutionary trajectories inferred from patient cohorts to reduce the number of solutions per patient [4, 14, 2, 17, 13]. Another approach that seeks to summarize the solution space is SubMARine, which infers a partial tree with confident ancestral relations [27]. These three classes of methods come with their own limitations. The sampling methods, which are typically MCMC-based, exhibit great bias to certain solutions [25], and thus may not infer a representative set of solutions. The consensus methods, including methods that utilize repeated evolutionary trajectories, require an exhaustive enumeration of all plausible trees, which is impractical to obtain when the set of possible trees is large.

To overcome these limitations, we introduce Sapling, a method that given read count data infers a small set of *backbone trees* on a smaller subset of mutations that collectively summarize the solution space (Fig. 1). We note that backbone trees are similar to the concept of a maximum-agreement subtree (MAST) in species phylogenetics [28], with a key distinction being that tumor phylogenies are node-labeled trees whereas species phylogenies are leaf-labeled trees. Using simulations, we show that the backbone trees returned by Sapling provide a good summary of the possible trees and can be expanded into full trees that are of higher quality than current state-of-the-art tree inference methods [18, 19, 29]. Finally, we demonstrate how Sapling can be applied to comprehensively summarize non-small lung cancer solution spaces with a small number of backbone trees [15].

**Fig. 1.**
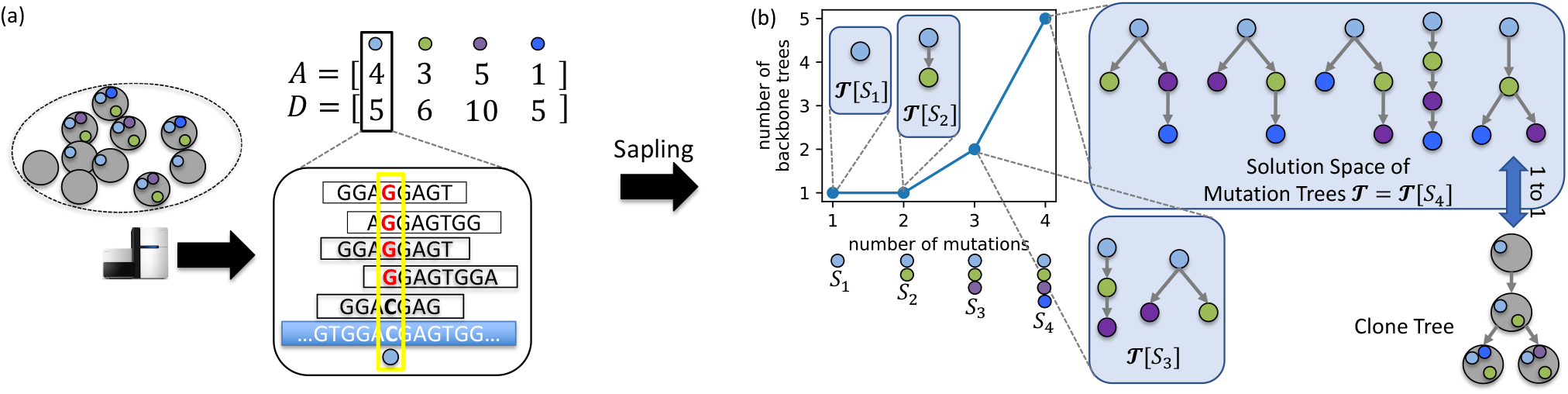
Overview of Sapling. (a) Bulk DNA sequencing, alignment and SNV calling results in matrices *A* and *D* of variant and total read counts of *n* SNVs in *m* samples. (b) Sapling is a heuristic for the Backbone tree inference from Reads problem, returning a small set of backbone trees for a given number *ℓ* of mutations. Here, with *ℓ* = 3 mutations, the solution space 𝒯 of 5 mutation trees can be summarized with two backbone trees 𝒯 [*S*_3_].

### Problem Statement

Due to uncertainty, cancer phylogeny inference algorithms typically infer a set 𝒯 of mutation trees from bulk sequencing data rather than a single tree. In this work, we consider trees inferred under the infinite sites assumption (ISA), meaning that each mutation is gained exactly once and never subsequently lost. We note that while this assumption does not generally hold particularly due to copy-number loss, tumor phylogeny pipelines include mutation clustering correcting for such events, yielding clusters of mutations that adhere to the ISA. Under the ISA, the solution space 𝒯 consists of rooted trees *T* whose nodes *V* (*T*) are labeled by mutations [*n*] = {1, …, *n*} — in practice, we will view mutation clusters as individual mutations. As such, we refer to nodes and mutations interchangeably. We write *u* ⪯_*T*_ *v* if node or mutation *u* occurs on the unique path from the root *r*(*T*) to node *v* — note that ⪯_*T*_ is reflexive, i.e., it holds that *u* ⪯_*T*_ *u* for mutations *u*. We denote the set of children of a node *v* of tree *T* by *δ*_*T*_ (*v*). We denote the parent of a node *v*≠ *r*(*T*) by *π*_*T*_ (*v*). Our goal is to identify common features or backbone trees on a smaller set *S* ⊆ [*n*] of mutations that best characterize the diversity of the solution space 𝒯. To that end, we define backbone trees as follows.

#### Definition 1

A rooted tree *T* [*S*] is a *backbone tree* of a tree *T* on mutations *S* ⊆ *V* (*T*) provided *u* ⪯_*T*_ *v* if and only if *u* ⪯_*T* [*S*]_ *v* for all mutations *u, v* ∈ *S*.

Mathematically, *T* is a subdivision or expansion of *T* [*S*] such that the backbone tree *T* [*S*] is obtained from *T* by contracting nodes *V* (*T*) \ *S*. Rather than considering a single backbone tree on a subset *S* of mutations, we wish to identify a backbone tree set 𝒯 [*S*] that collectively forms backbone trees of all trees 𝒯 on the full mutation set [*n*].

#### Definition 2

Given a set 𝒯 of trees on *n* mutations, the corresponding *backbone tree set* 𝒯 [*S*] for a subset *S* ⊆ [*n*] of mutations consists of all backbone trees *T* [*S*] of all trees *T* ∈ 𝒯.

Importantly, |𝒯 [*S*]| ≤ |𝒯 | for all *S* ⊆ [*n*]. The key question is which subset *S* ⊆ [*n*] of mutations provides an accurate summary of 𝒯 ? Ideally, we wish to simultaneously include as many mutations as possible in *S*, i.e. maximize |*S*|, while also minimizing the number of backbone trees |𝒯 [*S*]|. However, there is a tradeoff between both criteria. One can set *S* = [*n*], thus maximizing |*S*|, but this would lead to as many backbone trees as there are input trees, i.e. 𝒯 [*S*] = 𝒯, which does not provide a summary of 𝒯. On the other hand, setting *S* to contain no mutations or just a single mutation would lead to a backbone tree set consisting of a single backbone tree composed of at most one mutation; thus while minimizing |𝒯 [*S*]| = 1, this does not provide any useful information that is particular to 𝒯. To model this tradeoff, we formulate the following two problem statements, constraining either the number |*S*| of mutations is constrained or the number |𝒯 [*S*]| of backbone trees.

#### Problem 1

(backbone Tree Set Inference) Given a set 𝒯 of trees on *n* mutations and parameter *ℓ* ∈ [*n*], find a subset *S* ⊆ [*n*] of *ℓ* mutations and corresponding backbone tree set 𝒯 [*S*] such that 𝒯 [*S*] has minimum cardinality among all backbone tree sets induced by *ℓ* mutations.

#### Problem 2

(backbone Tree Set Inference) Given a set 𝒯 of trees on *n* mutations and parameter *τ* ∈ N, find a maximum-cardinality subset *S* ⊆ [*n*] of mutations and corresponding backbone tree set 𝒯 [*S*] such that |𝒯 [*S*]| ≤ *τ*.

A special version of this problem arises when *τ* = 1. In that case, we are seeking a maximum-cardinality set *S* of mutations and a single corresponding backbone tree *T* [*S*] on which all trees in 𝒯 agree. In our final problem, we seek to expand a given backbone tree into a full tree.

#### Problem 3

(backbone Tree Expansion) Given trees 𝒯 on *n* mutations and a tree *T* [*S*] on a subset *S* ⊆ [*n*] of mutations, find a tree *T* ∈ 𝒯 such that *T* [*S*] is backbone tree of *T*.

### Backbone Tree Inference and Expansion from Reads

In practice, we are not given the set 𝒯 of phylogenetic trees. While there exist many methods for inferring such a set via sampling [5, 19, 29, 18] or enumeration [21, 8], obtaining the complete set 𝒯 of trees might be infeasible due to its sheer size [25]. Therefore, we propose to infer backbone trees directly from read count data obtained from bulk DNA sequencing of *m* samples (regional or temporal) from the same tumor. More specifically, we are given two matrices *A, D* ∈ ℕ ^*m*×*n*^ where *A* = [*a*_*p,i*_] indicate the number of reads supporting the variant allele and *D* = [*d*_*p,i*_] the total number of reads at each mutation locus in each sample.

To pose the problem, we are interested in computing the probability Pr(*T* | *V, D*) of a tree *T* given the data *A, D*. To compute this probability, we note that the number *a*_*p,i*_ of variant read counts at mutation locus *i* in sample *p* depends on the total number *d*_*p,i*_ of reads, i.e. 0 ≤ *a*_*p,i*_ ≤ *d*_*p,i*_, and the (latent) frequency *f*_*p,i*_ ∈ [0, 1] of mutation *i* in sample *p*. Typically, one models variant read counts *A* = [*a*_*p,i*_] as binomial distributions, i.e. *a*_*p,i*_ ∼ binom(*d*_*p,i*_, *f*_*p,i*_). Using the independence of mutations and samples, we thus have

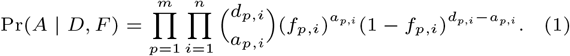

As discussed in [7], frequencies *F* depend on a tree *T*. That is, a tree *T* under the ISA constrains frequencies *F* = [*f*_*p,i*_] as

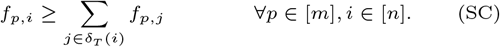

This is also known as the sum condition (SC). To compute the desired probability Pr(*T* | *V, D*), we apply Bayes’ rule, yielding Pr(*T* | *V, D*) = [Pr(*A, D* | *T*) Pr(*T*)]*/* Pr(*A, D*), which is proportional to Pr(*A, D* | *T*) using a flat prior on Pr(*T*). We now have

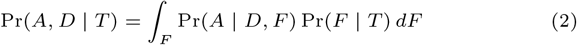

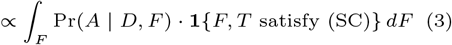

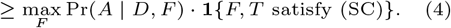

In other words, we approximate the probability Pr(*A, D* | *T*) of read counts *A, D* given a tree *T* by seeking a frequency matrix *F* such that *F* and *T* satisfy (SC) and Pr(*A* | *D, F*) is maximum. This is equivalent to solving the following optimization problem.

#### Definition 3

Given reads *A, D*, the *log-likelihood* ℒ(*A, D* | *T*) of a rooted tree *T* equals max_*F*_ ℒ(*A, D* | *F*) s.t. (SC) where ℒ(*A, D* | *F*) is defined as 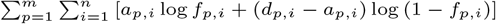.

Computing ℒ(*A, D* | *T*), or equivalently −ℒ(*A, D* | *T*), requires solving a convex optimization problem subject to linear constraints, which can be solved in polynomial time (for a fixed error tolerance) using interior point methods [22]. To allow one to obtain near-maximum likelihood solutions, the user may specify the parameter *ρ* ∈ [0, 1] yielding the solution space 𝒯 ^(*ρ*)^ of trees that are most a factor of *ρ* removed from maximum likelihood, formally defined as follows.

#### Definition 4

Given *ρ* ∈ [0, 1] and read counts *A, D* ∈ ℕ ^*m*×*n*^, the set 𝒯 ^(*ρ*)^ includes all trees *T* such that Pr(*A, D* | *T*) ≥ *ρ* Pr(*A, D* | *T* ^*^) where *T* ^*^ is a tree with maximum probability.

Thus, we have 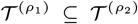 for all 0 ≤ *ρ*_1_ ≤ *ρ*_2_ ≤ 1. Specifically, for *ρ* = 0 the set 𝒯 ^(0)^ contains all *n*^*n*−1^ rooted trees on *n* mutations [3]. This leads to the following updated problem statements.

#### Problem 4

(backbone Tree Inference from Reads) Given variant and total read counts *A, D* ∈ ℕ ^*m*×*n*^ for *n* mutations in *m* samples and parameters *ℓ* ∈ [*n*] and *ρ* ∈ [0, 1], find a subset *S* of *ℓ* mutations and corresponding backbone tree set 𝒯 [*S*] such that 𝒯 [*S*] has minimum cardinality among all backbone tree sets induced by *ℓ* mutations on trees 𝒯 that are at most a factor of *ρ* removed from maximum likelihood.

#### Problem 5

(backbone Tree Inference from Reads)(Backbone Tree Inference from Reads) Given variant and total read counts *A, D* ∈ ℕ ^*m*×*n*^ for *n* mutations in *m* samples and parameters *τ* ∈ ℕ and *ρ* ∈ [0, 1], find a maximum-cardinality subset *S* ⊆ [*n*] of mutations and backbone tree set 𝒯^(*ρ*)^ [*S*] such that |𝒯^(*ρ*)^ [*S*]| ≤ *τ*.

#### Problem 6

(Backbone Tree Expansion from Reads) Given variant and total read counts *A, D* ∈ N^*m*×*n*^ for *n* mutations in *m* samples and a tree *T* [*S*] on a subset *S* ⊆ [*n*] of mutations, find a tree *T* ∈ 𝒯^(*ρ*)^ such that *T* [*S*] is a backbone tree of *T*.

## Methods

In this section, we introduce the two algorithms that make up Sapling. First, we introduce a heuristic to solve the Backbone Tree Inference from Reads problem subject to either a constraint on the number of mutations or the number of backbone trees. Second, we introduce a heuristic to solve the Backbone Tree Expansion from Reads problem.

### Enumerating backbone trees

Given *A* = [*a*_*p,i*_] and *D* = [*d*_*p,i*_] it is clear that 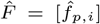 where 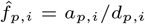 is the frequency matrix that maximizes ℒ(*A, D* | *T*) when ignoring the sum condition (SC). The question whether there exists a tree *T* satisfying (SC) for a given a frequency matrix was shown to NP-complete when *F* contains *m* ≥ 2 samples [8]. This means that the Backbone Tree Inference from Reads problem from reads is NP-hard when *m* ≥ 2. To see why, we can set *ρ* = 1 and solve the problem with *ℓ* = *n* or *τ* = *n*^*n*−1^. This will return one or more trees on all *n* mutations. If the likelihood ℒ(*A, D* | *T*) of any such tree *T* equals the likelihood 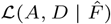 then there exists a tree satisify the sum condition with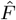, leading to the following hardness result.

#### Theorem 1

*The two* Backbone Tree Inference from Reads *problems are NP-hard even when m* = 2.

*Proof* We show hardness by a polynomial-time reduction from the Perfect Phylogeny Mixture Deconvolution (PPMD) problem [7, 8, 25] of deciding whether there exists a tree *T* satisfying (SC) for a given frequency matrix *F* ∈ [0, 1]^*m*×*n*^. This decision problem was shown to NP-complete when *F* contains *m* ≥ 2 samples [8]. Without loss of generality, we may assume that *F* is rational (the reduction from Subset Sum presented in [8] works for rational values). Thus, we have *f*_*p,i*_ = *a*_*p,i*_*/d*_*p,i*_ where *a*_*p,i*_, *d*_*p,i*_ ∈ ℕ and *a*_*p,i*_ ≤ *d*_*p,i*_ for each sample *p* ∈ [*m*] and mutation *i* ∈ [*n*]. These entries correspond to variant read count matrix *A* = [*a*_*p,i*_] and total read count matrix *D* = [*d*_*p,i*_]. This reduction takes polynomial time.

We claim that Problem 4 with read counts *A, D* obtained from *F* is NP-hard when *m* ≥ 2, *ρ* = 1 and *ℓ* = *n*. Moreover, we claim that Problem 5 with read counts *A, D* obtained from *F* is NP-hard when *m* ≥ 2, *ρ* = 1 and *τ* = *n*^*n*−1^. Solving either problem will return a set 𝒯 of trees. Let *T* ∈ 𝒯 be one such solution tree. Clearly, *T* is a tree on *n* mutations due to the constraint *ℓ* = *n* and *τ* = *n*^*n*−1^. Since *F* is the maximum likelihood estimator of the binomial proportions of *A, D*, we have that ℒ(*A, D* | *T*) ≤ ℒ(*A, D* | *F*). Furthermore, the likelihood ℒ(*A, D* | *T*) equals the likelihood ℒ(*A, D* | *F*) if and only if *T* satisfies (SC). That is, we can verify whether this bound is tight by simply checking whether *F, T* satisify (SC), which takes polynomial time. □

We note that the second version of Backbone Tree Inference from Reads problem where we are given an upperbound *τ* of backbone trees can be solved by repeatedely solving the variant of the problem where the number *ℓ* of mutations is fixed. That is, starting with *ℓ* = 1, we obtain the backbone tree set 𝒯_*ℓ*_ and increment *ℓ* until the resulting number |𝒯_*ℓ*_| of trees is greater than *τ*. We then return |𝒯_(*ℓ*−1)_|. Since the Backbone tree inference from Reads problems are hard, we introduce the following heuristic.

#### A naive approach

In our first approach, we propose to build the backbone trees iteratively. We intialize the set 𝒯_1_ with a single tree *T* containing a single mutation (can be any mutation). Then at each iteration *k* ≥ 1, given the current set 𝒯_*k*_ on the same subset *S*_*k*_ of mutations, we extend each tree *T* ∈ 𝒯_*k*_ by adding a new mutation *i* ∈ [*n*] \ *V* (*T*) at each possible location. Specifically, we may either extend *T* by adding the edge (*i, r*(*T*)), or for an existing node *j* ∈ *V* (*T*) we insert the new edge (*j, i*) and distribute the original children *δ*_*T*_ (*i*) among nodes *i* and *j*. This results in a new set 𝒯_*k*+1_ of trees on mutations *S*_*k*_ ∪ {*i*}.

In order to assess whether an expanded tree *T* ^′^ ∈ 𝒯_*k*+1_ is a good backbone tree on the mutation set *S*_*k*+1_ = *S*_*k*_ ∪ {*i*}, we evaluate the likelihood ℒ(*A*[*S*_*k*+1_], *D*[*S*_*k*+1_] | *T* ^′^). That is, we take the submatrices *A*[*S*_*k*+1_] of *A* and *D*[*S*_*k*+1_] of *D* with columns corresponding to mutations *S*_*k*+1_. Computing this likelihood requires solving a convex optimization problem and, as mentioned before, this can be solved efficiently in polynomial time with a constant error tolerance. We calculate ℒ(*A*[*S*_*k*+1_], *D*[*S*_*k*+1_] | *T* ^′^) for all trees *T* ^′^ in 𝒯_*k*+1_, retaining only those trees that have a likelihood of at least *ρ* · Pr(*A*[*S*_*k*+1_], *D*[*S*_*k*+1_] | *T* ^*^), where *T* ^*^ is the maximum likelihood tree among 𝒯_*k*+1_. We terminate after iteration *ℓ* − 1, returning the set *S*_*ℓ*_ of mutations and backbone tree set 𝒯_*ℓ*_.

However, this algorithm does not work well in practice for two key reasons. First, the order in which mutations are considered does affect the number of resulting backbone trees. One might overcome this limitation by exploring different permutations of mutations, but this becomes quickly intractable as there are *n*! permutations. Ideally, one would be able to determine a good permutation of mutations ahead of time, and only consider this single permutation when enumerating. Second, note there are 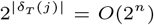 possible expanded trees *T* ^′^ when expanding a single mutation *j* of a partial tree *T*. This would make it impossible to explore the entire set of possible backbone trees at each iteration. Therefore, we need additional criteria to prune the search space.

#### Adding mutations ordered by 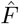

While we do not know the true latent frequencies of the mutations, 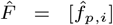 where 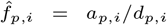 can serve as a good estimator of the latent frequencies. As such, we propose to sort the *n* mutations in descending order based on 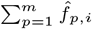 and consider the mutations according to this order when enumerating backbone trees. Intuitively, this order will start by adding mutations that are closest to the root and leave the mutations that are farthest away as the last mutations to be added. We note that Orchard uses this same ordering when sampling complete trees from the solution space [18].

#### Pruning the search space

To avoid considering *O*(2^*n*^) possible trees when expanding a partial tree *T*, we propose to place a new mutation *i* either (i) as the new root of *T*, or (ii) as a new leaf of *T*, or (iii) split an existing edge (*π*_*T*_ (*j*), *j*) of *T* inserting edges (*π*_*T*_ (*j*), *i*) and (*i, j*). Importantly, there are only *O*(*n*) such possible expansions of a given tree *T*. There is a theoretical justification for case (ii) of this pruning step when the *n* mutations are sorted such that 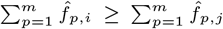 for any 1 ≤ *i < j* ≤ *n*. If the sequencing depth is large enough, 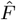 accurately reflects the latent frequencies of the mutations — i.e. 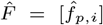 is an unbiased maximum likelihood estimator if the variants read count are indeed binomially distributed. In this case, for any *ρ >* 0, each tree *T* ∈ 𝒯 ^(*ρ*)^ must adhere to (SC) with respect to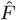. As such, we have that a new mutation *j > i* cannot be a parent of a previous mutation *i*.

##### Theorem 2

*Let* 0 *< ρ* ≤ 1 *and let* 𝒯 ^(*ρ*)^ *be the set of corresponding trees on n mutations. As d*_*p,i*_ → ∞ *for all p and i, then if* 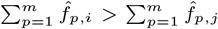, *it holds that j ⊀*_*T*_ *i for all trees T* ∈ 𝒯 ^(*ρ*)^.

*Proof* We prove this by contradiction, assuming there exists a tree *T* ∈ 𝒯 ^(*ρ*)^ with mutations *i, j* such that mutation *j* is ancestral to mutation *i* while 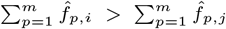. Note that as *d*_*p,i*_ → ∞ for all *p* and *i*, the probabilty distribution of *v*_*p,i*_ is concentrated at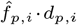. Therefore, if *T* does not adhere to (SC) w.r.t. 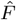 then ℒ(*A, D* | *T*) → −∞. This in turn would imply that *T* is not in 𝒯 ^(*ρ*)^ for any *ρ >* 0. Hence, *T* adheres to (SC) w.r.t.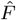. As such, we have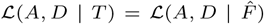. By (SC), we have 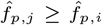 for any sample *p* ∈ [*m*]. Hence,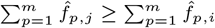, which contradicts the assumption. □

In practice, entries *D* = [*d*_*p,i*_] do not go to infinity. Therefore, we consider additional inclusion criteria beyond adding mutation *i* as a new leaf of *T*. Specifically, we allow a new mutation *i* to be a parent of an existing mutation *j* (cases (i) and (iii)). Since we do not expect the real *F* to deviate too much from 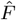, it is unlikely that mutation *i* will be assigned more than one child of mutation *j* in tree *T*, allowing us to avoid enumerating all possible redistributions of the original children of *j* among *i* and *j*. This in turn limits the number of maximum likelihood calculations. The pseudocode of the updated procedure FastBackboneEnumeration is given in Algorithm 1, where AddMutationFast(*T, i*) corresponds to the faster method of adding mutation *i* to tree *T* discussed above.

##### Algorithm 1

FastBackoneEnumeration(*A,D,ℓ,ρ*)

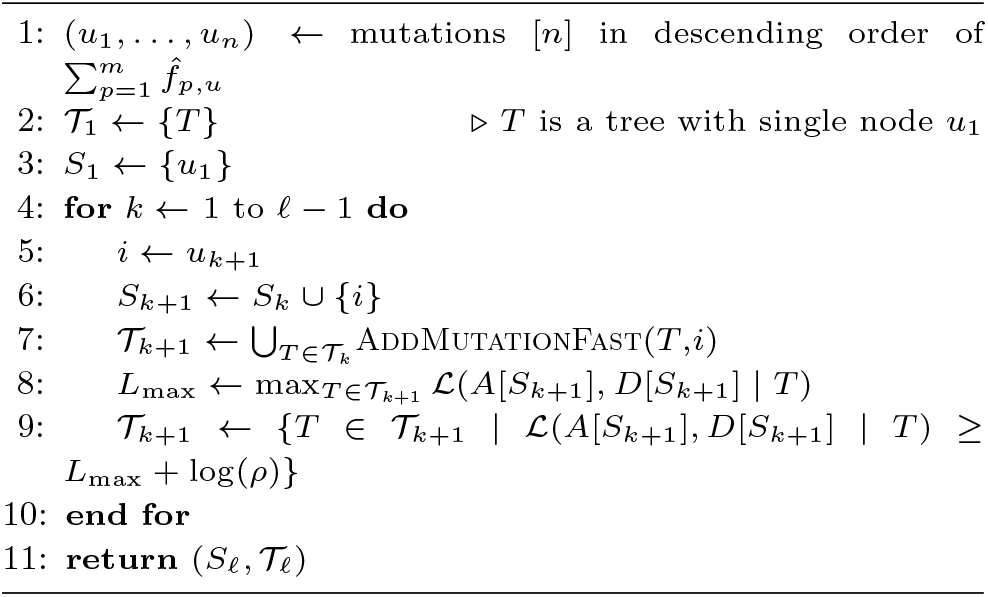

### Expanding a backbone tree into a full tree

Note that solving Backbone Tree Expansion from Reads given an empty tree is equivalent to finding a maximum likelihood tree given reads *A, D*, which is NP-hard as discussed. In the following, we propose a greedy algorithm to solve this problem heuristically. We employ similar ideas of growing the tree as in the FastBackboneEnumeration algorithm, considering the remaining mutations [*n*] \ *V* (*T*) in descending order according to 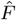. However, rather than keeping a large set of trees that have a high likelihood, we only retain a single tree with the highest likelihood at each iteration. The pseudocode of this method called GreedyExpansion is given in Algorithm 2.

#### Algorithm 2

GreedyExpansion(*A,D,T*)

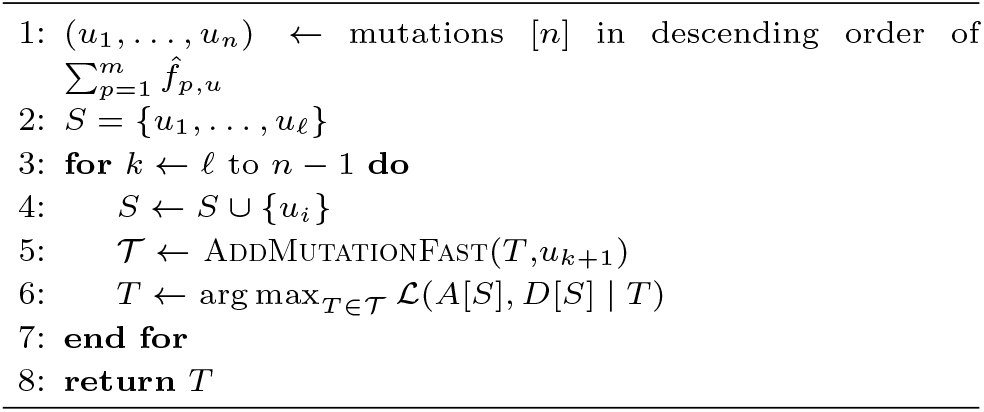

### Implementation details

Sapling provides a Python implementation of the FastBackboneEnumeration and GreedyExpansion algorithms. As mentioned above, computing −ℒ(*A, D* | *T*) is equivalent to solving a convex optimization problem with a convex objective function subject to linear constraints. Therefore, Sapling use the Python package `cvxopt` to optimize −ℒ(*A, D* | *T*) efficiently. In line 8 of FastBackoneEnumeration (Algorithm 1), Sapling uses an error tolerance to account for floating point errors and the possibility of underestimating the likelihood. If clusters of mutations are provided rather than individual mutations, Sapling takes the median depth of all mutations in the cluster as the depth of the cluster and the median depth times the average frequency 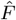 as the variant reads (rounded to the nearest integer) for the cluster and treat the cluster as a single mutation. Sapling is available at https://github.com/elkebir-group/Sapling.git under the 3-Clause BSD open source license.

## Results

In this section, we evaluate Sapling on (i) a set of small simulation instances with known optimal solutions, (ii) a set of larger simulation instances and (iii) real data. Specifically, we compare Sapling’s backbone trees (obtained via FastBackboneEnumeration) to the optimal backbone tree sets and to alternative summarization obtained by the MCT algorithm [1]. Moreover, we compare Sapling’s expanded trees (obtained via GreedyExpansion) to trees sampled by Pairtree [19, 29] and Orchard [18].

### Simulation setup

To generate a simulation instance with a number *n* of mutations and number *m* of samples, we start by randomly generating an unrooted, node-labeled tree *T* with *n* nodes/mutations using Prüfer sequences [24]. Next, we root the tree *T* uniformly at random, followed by drawing *m* samples of fractions of the *n* clones corresponding to the nodes in the tree from a Dirichlet distribution, ultimately yielding frequency matrix *F* = [*f*_*p,i*_]. For each frequency *f*_*p,i*_, the total number *d*_*p,i*_ of reads is drawn from a Poisson distribution with mean *λ* = 100, simulating an average sequencing depth of 100×. Finally, the variant reads *A* = [*a*_*p,i*_] are each drawn from a binomial distribution with *d*_*p,i*_ trials and success probability *f*_*p,i*_.

### Sapling identifies near-optimal backbone trees

We generated a simulation dataset of 20 instances with *n* = 8 mutations and *m* = 2 samples. The small number *n* = 8 of mutations allowed us to exhaustively enumerate the 𝒯 ^(0.9)^ of complete trees with a likelihood that is at most a fraction of *ρ* = 0.9 away from maximum likelihood. To accomplish this, we generated all *n*^*n*−1^ = 8^7^ = 2,097,152 trees *T* and computed their likelihoods ℒ(*A, D* | *T*). We ran Sapling’s FastBackboneEnumeration algorithm with parameters *ℓ* ∈ {1, …, 8} as well as *τ* ∈ {1, 2, 5, 10, 20, 50}.

We show the number |𝒯 ^(0.9)^| of trees in Fig. 2a, ranging from 3 to 672 with a median of 65 trees. Next, we enumerated all 2^8^ = 256 subsets *S*^′^ ⊆ [*n*] of mutations, and identified the number |𝒯 ^(0.9)^[*S*^′^]| of backbone trees for each subset *S*^′^ of mutations. This allowed us to compare the number |𝒯 ^(0.9)^[*S*]| of backbone trees returned by Sapling for varying values of *ℓ* to the optimal number |𝒯 ^(0.9)^[*S*^*^]| of backbone trees such that |*S*^*^| = *ℓ* by computing the *approximation ratio* defined as |𝒯 ^(0.9)^[*S*]|*/*|𝒯 ^(0.9)^[*S*^*^]|. Thus, an approximation ratio of 1 indicates that Sapling identified an optimal (minimum) set of backbone trees. We find the median approximation ratio is 1 with a maximum ratio of 7 with |𝒯 ^(0.9)^[*S*]| = 14 inferred backbone trees by Sapling versus |𝒯 ^(0.9)^[*S*^*^]| = 2 optimal backbone trees for *ℓ* = 5 mutations (Fig. 2b). In Fig. 2c, we show an instance where Sapling returned optimal solutions for all *ℓ*, whereas Fig. 2d shows an instance where Sapling did not return optimal solutions for all *ℓ*. Specifically, for *ℓ* = 3 Sapling returned two backbone trees for mutations *S* = {0, 1, 3} shown in Fig. 2e whereas there exists a different set *S*^*^ = {0, 3, 4} with just a single backbone tree shown in Fig. 2f.

**Fig. 2.**
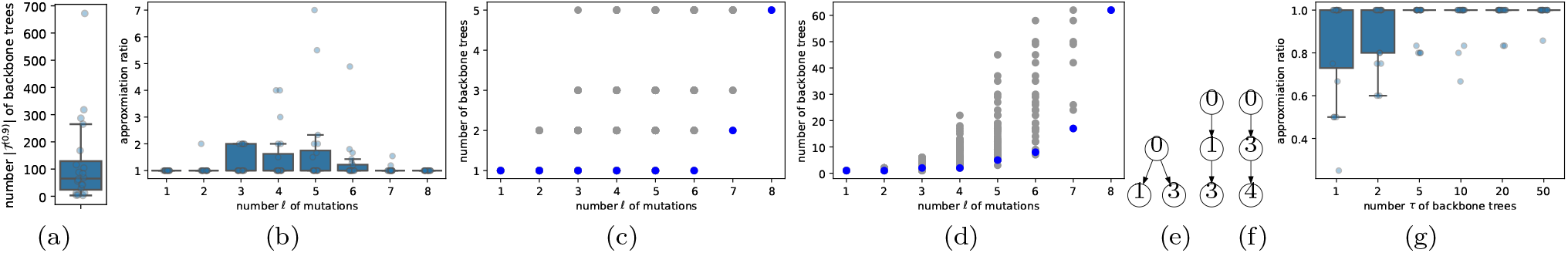
Simulations on *n* = 8 mutations and *m* = 2 samples. (a) Set 𝒯 ^(0.9)^ of trees that have a likelihood that is at most a factor of *ρ* = 0.9 away from maximum likelihood. (b) Approximation ratio achieved by Sapling for varying *ℓ*. (c) A simulation instance that Sapling (blue) solves to optimality for all *ℓ*, with gray entries indicating subsets of mutations not considered by Sapling. (d) A simulation instance that Sapling did not solve to optimality for *ℓ* ∈ {3, 5, 6}. (e) The two bacbone trees retured by Sapling for the instance shown in (d) at *ℓ* = 3. (f) The optimal backbone tree of the instance in (e) determined by exhaustive enumeration at *ℓ* = 3. (g) Approximation ratio achieved by Sapling for varying *τ*.

Similarly, we evaluate the approximation ratio when running Sapling with a specified upperbound *τ* of backbone trees. Specifically, let |*S*| be the number of mutations returned by Sapling and |*S*^*^| be the maximum number of mutations such that |𝒯 ^(0.9)^[*S*]|, |𝒯 ^(0.9)^[*S*^*^]| ≤ *τ*. The *approximation ratio* equals |*S*|*/*|*S*^*^|, where a value of 1 indicates that Sapling returned the optimal (maximum) number of mutations and a value smaller than 1 indicates that Sapling underestimated the number of mutations. Again, we find that the median approximation ratio is 1, with a minimum ratio of 0.25 for which Sapling identified |*S*| = 1 mutations versus a maximum number |*S*^*^| = 4 of mutations for *τ* = 1 (Fig. 2c). In particular, for smaller *τ* the approximation ratio may be smaller than 1.

Thus, in general we find that the heuristic employed by Sapling in the majority of cases finds an optimal solution for these small simulation instances. Moreover, the backbone trees returned by Sapling have perfect recall compared to the backbone tree set obtained from the ground-truth complete trees using the same set of mutations (data not shown).

### Sapling infers high-quality backbone and full trees

In this section, we demonstrate Sapling is also capable of handling larger input instances. To that end, we generated 60 additional simulation instances with *m* = 10 samples and *n* ∈ {20, 50, 100} mutations (with 20 instances for each value of *n*). We ran Sapling on a laptop with 16 GB RAM and an Apple M1 Pro CPU. We show the running time of Sapling’s backbone tree enumeration mode in Fig. 3a, showing an exponential increase in running time with increasing number *n* of mutations and increasing values of the parameter *τ* ∈ {1, 5, 10, 20, 50}.

**Fig. 3.**
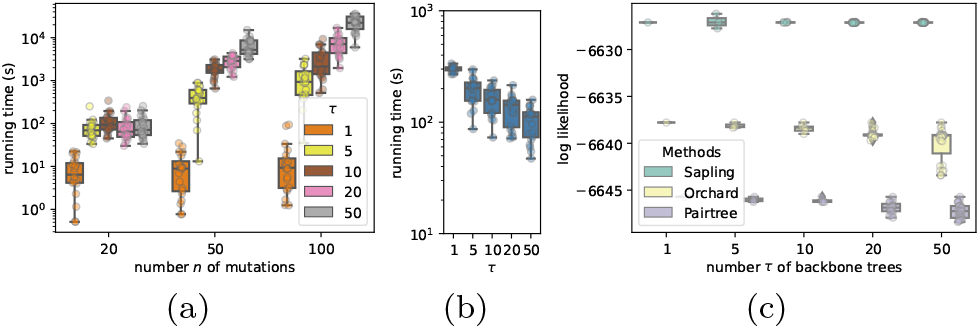
Simulations on *n* ∈ {20, 50, 100} mutations and *m* = 10 samples. (a) Running time of Sapling’s backbone tree enumeration algorithm for varying values of *τ*. (b) Running time of Sapling’s backbone tree expansion algorithm when given an initial backbone tree obtained using the specified *τ* parameter for simulation instances with *n* = 50 mutations. (c) Likelihood values of complete trees identified by Sapling, Orchard and Pairtree for a single *n* = 50 simulation instance.

We additionally ran Sapling’s backbone tree expansion algorithm to expand each identified backbone tree into a full tree for all twenty *n* = 50 simulation instances. We find that the running time ranged from a minimum of 47 seconds when provided a *τ* = 50 backbone tree (containing 47 mutations) vs. 335 seconds when provided a *τ* = 1 backbone tree (containing 12 mutations) — see Fig. 3b.

To compare Sapling’s complete trees, we also ran Pairtree and Orchard and retained their *τ* highest likelihood unique trees (we ran these algorithms with default parameters using 4 MCMC chains with 2500 samples each and a burn-in of 1250 samples for Pairtree; and a beam width of *k* = 10, a branching factor of *f* = 100 and 8 parallel instances for Orchard). The comparison of the log-likelihood of one *n* = 50 simulation instance is shown in Fig. 3c, showing that Sapling’s trees achieved higher likelihoods (median: −6627.12 for *τ* = 50) than those identified by Pairtree (median: −6647.23 for *τ* = 50) and Orchard (median: −6639.16 for *τ* = 50). It is important to note, however, that Pairtree and Orchard (79 × 4 and 59 × 8 seconds, respectively) ran much faster than Sapling (2 hours and 30 minutes for backbone enumeration and 31 minutes for backbone expansion for *τ* = 50).

In summary, Sapling can be used to obtain a diverse set of high-likelihood trees by expanding initial backbone trees.

### Sapling summarizes the solution space of real data

Finally, we ran Sapling on the TRACERx cohort of 100 non-small-cell lung cancer patients [15] using the mutation clusters reported by the authors, whose number *n* of clusters ranged from 2 to 15 and number *m* of sequencing samples ranged from 1 to 7. For each patient, we ran Sapling with parameters *ρ* ∈ {0.4, 0.9} and *ℓ* ∈ {1, …, *n*}. Sapling’s running time ranged from less than 1 second to 90 seconds (Apple M1 Pro CPU with 16 GB RAM, data not shown).

Setting *ℓ* = *n* results in Sapling enumerating the complete solution space 𝒯 ^(*ρ*)^ for the specified value of *ρ*, indicating the allowed deviation from maximum-likelihood. The distribution of 𝒯 ^(*ρ*)^ is shown in Fig. 4a, showing that the number of trees increased with decreasing *τ* as expected. Specifically, there are 26 and 14 patients with at least two trees in 𝒯 ^(*ρ*)^ for *ρ* = 0.4 and *ρ* = 0.9, respectively.

**Fig. 4.**
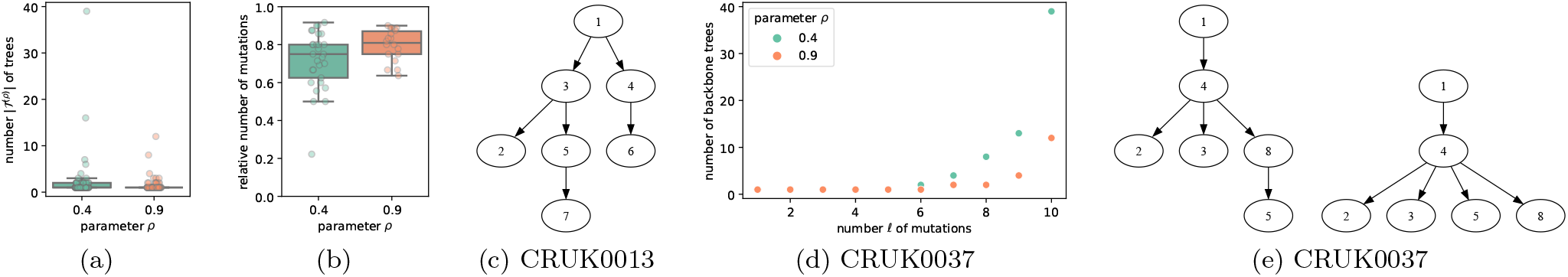
Sapling performance on TRACERx data. (a) The number |𝒯 ^(*ρ*)^ | of complete trees identified by Sapling for *ρ* ∈ {0.4, 0.9}. (b) The fraction of mutation clusters in the backbone tree for *τ* = 1 (only showing patients where |𝒯 ^(*ρ*)^ | *>* 1). (c) The single backbone tree identified by Sapling with *τ* = 1, *ρ* = 0.4 for patient CRUK0013. (d) The number of backbone trees identified by Sapling for varying number *ℓ* of mutations for patient CRUK0037. (e) The two backbone trees identified by Sapling with *τ* = 2, *ρ* = 0.4 for patient CRUK0037.

On the other hand, when setting *τ* = 1, Sapling seeks to identify a single backbone tree with maximum number of mutations. In Fig. 4b, we show the fraction of mutations that are included in each individual backbone tree per patient, finding that a median fraction of 0.75 and 0.81 of mutations is included for *ρ* = 0.4 and *ρ* = 0.9, respectively. We show the *τ* = 1 backbone tree identified by Sapling with *ρ* = 0.4 for patient CRUK0013 with *n* = 9 mutation clusters. This backbone tree spans 7 out of 9 mutation clusters, and is a proper subtree of the single consensus tree for CRUK0013 reported by the MCT algorithm [1]. For patient CRUK0037 with *n* = 10 mutation clusters, restricting the number of backbone trees to *τ* = 1 results in only 5 and 6 covered mutation clusters for *ρ* = 0.4 and *ρ* = 0.9, respectively (Fig. 4d). With *τ* = 2 backbone trees, Sapling covers an additional mutation cluster for both values of *ρ*. We show the two backbone trees for *ρ* = 0.4 in Fig. 4e, which, again, form proper subtrees of the two consensus trees reported by the MCT algorithm for this patient [1].

In summary, on real data, we find that Sapling is able to quickly enumerate the complete solution space of trees and comprehensively summarize it with a small number of backbone trees that span a large fraction of mutations.

## Discussion

In this work, we introduced the Backbone Tree Inference from Reads and Backbone Tree Expansion from Reads problems, which, respectively, seek to identify or expand a set of backbone trees given read count data. We showed that these problems are NP-hard, and introduced a heuristic algorithm, Sapling. Using simulations, we showed that Sapling provides a good approximate solution to both Backbone Tree Inference from Reads problems. We also demonstrated that Sapling returns more plausible trees than the current state-of-the-art methods, Pairtree [19, 29] and Orchard [18]. On real data, we ran Sapling on the TRACERx cohort of 100 lung cancer patients [15], showing that Sapling’s backbone trees adequately summarize the solution space of trees.

There are several future directions. First, we could relax the infinite sites assumption and support mutation loss. Second, given the decrease in performance for *τ* = 1, we plan on developing a specialized algorithm for this case. Third, we plan to sample from backbone trees rather expanding these greedily.

